# MS-CleanR: A feature-filtering approach to improve annotation rate in untargeted LC-MS based metabolomics

**DOI:** 10.1101/2020.04.09.033308

**Authors:** Ophélie Fraisier-Vannier, Justine Chervin, Guillaume Cabanac, Virginie Puech-Pages, Sylvie Fournier, Virginie Durand, Aurélien Amiel, Olivier André, Omar Abdelaziz Benamar, Bernard Dumas, Hiroshi Tsugawa, Guillaume Marti

**Author notes:** Corresponding author; mail.

## Abstract

Untargeted metabolomics using liquid chromatography-mass spectrometry (LC-MS) is currently the gold-standard technique to determine the full chemical diversity in biological samples. This approach still has many limitations, however; notably, the difficulty of estimating accurately the number of unique metabolites being profiled among the thousands of MS ion signals arising from chromatograms. Here, we describe a new workflow, MS-CleanR, based on the MS-DIAL/MS-FINDER suite, which tackles feature degeneracy and improves annotation rates. We show that implementation of MS-CleanR reduces the number of signals by nearly 80% while retaining 95% of unique metabolite features. Moreover, the annotation results from MS-FINDER can be ranked with respect to database chosen by the user, which improves identification accuracy. Application of MS-CleanR to the analysis of *Arabidopsis thaliana* grown in three different conditions improved class separation resulting from multivariate data analysis and lead to annotation of 75% of the final features. The full workflow was applied to metabolomic profiles from three strains of the leguminous plant *Medicago truncatula* that have different susceptibilities to the oomycete pathogen *Aphanomyces euteiches*; a group of glycosylated triterpenoids overrepresented in resistant lines were identified as candidate compounds conferring pathogen resistance. MS-CleanR is implemented through a Shiny interface for intuitive use by end-users (available at: https://github.com/eMetaboHUB/MS-CleanR).

---

Untargeted, or discovery-based metabolomics has become an essential tool in all biological sciences including clinical research^1,2^, plant science^3^ and natural product mining^4^, among many other applications. Living organisms are estimated to contain more than one million distinct compounds^5^. According to the MetaboLights database (DB), 80% of untargeted metabolomics workflows rely on liquid chromatography-mass spectrometry (LC-MS) (https://www.ebi.ac.uk/metabolights/). Due to its broad coverage of metabolites, LC-MS based metabolomics has become the preferred tool to detect several hundreds of compounds encountered in a complex biological material. Many software programs have been developed to turn features (*m/z* × retention time (RT) pairs) extracted from LC-MS raw data into chromatographic peak lists, including web-based interfaces such as XCMS^6^, Workflow4Metabolomics^7^, local GUI with MZmine^8^ and MS-DIAL^9^. Despite significant progress in feature extraction, it remains a challenge to estimate accurately the number of unique metabolites in a crude extract from the profile of one LC-MS experiment^10^. On average, untargeted LC-MS profiling yields hundred to thousands of features, which include isotopes, contaminants, adducts, dimers, multimers and heteromeric complexes, and artifacts. Patti and colleagues^11^ used the term ‘degenerate features’ to describe multiple signals derived from the same metabolite; they demonstrated that feature inflation is highly underestimated and insufficiently addressed in untargeted LC-MS based metabolomics. This may have important consequences by increasing both the false annotation rate and the number of ‘unknown’ features arising from wrongly attributed signals. This is especially true when the annotation process is based on *in silico* modeling of fragmentation patterns, as are Sirius^12^, MS-FINDER^13^, MetFrag^14^ or CFM-ID^15^, since tandem mass spectrometry (MS/MS) spectra are processed without taking into account feature relationships. Thus, most untargeted metabolomics studies focus on a subset of identified metabolites for which spectral data are easily accessible from public repositories or in-house DBs.

A few packages have been developed to deal with feature degeneracy: CAMERA^16^ is based on adduct relationships; RAMClust^17^ correlates features in multiple samples; MS-FLO^18^ uses Pearson’s correlation and peak height similarity to identify adducts, duplicate peaks and isotope features of the main monoisotopic ion, and MZunity^19^ which confronts adducts or neutral loss index to decipher relationship among the acquired high resolution pseudo-molecular ions list. Deep-learning approaches have also been developed based on LC-MS spectral peak shape filtering^20,21^. All these packages focus on a single type of degeneracy, however, and they are difficult to implement in a unified workflow.

Among the most advanced and versatile methods developed recently for untargeted metabolomics is the tandem MS-DIAL-MS-FINDER suite. MS-DIAL is an all-in-one program for metabolomics and lipidomics that relies on mass spectral libraries such as NIST 14 and MassBank of North America (MoNA) for metabolite annotation. MS-FINDER is a partner program of MS-DIAL, in which unknown structures can be elucidated from MS/MS spectra by the hydrogen rearrangement rules-based scoring system. Here, we describe a third tool in this suite, called MS-CleanR, to remove degenerate features and improve annotation rates from untargeted LC-MS-based metabolomics data. Starting from the aligned peak list files determined by the MS-DIAL deconvolution process, our R package firstly removes noise signals by using generic filters. In the second step, the package groups the ion features based on the results of the MS-DIAL peak character estimation algorithm^22^ providing the ion linkages of adducts, correlated chromatograms, putative ion source fragments candidates and similar metabolite profiles among samples. In the third step, clustered ion features are merged between positive ionization (PI) and negative ionization (NI) modes and the adduct relationships are corrected accordingly. The cleaned-up feature list can be exported to MS-FINDER for annotation purposes. Finally, the package merges the MS-FINDER annotation output with the cleaned-up peak list and includes the possibility to prioritize identification according to the DBs used for MS-FINDER interrogation. The whole MS-CleanR workflow is easily accessible through a Shiny user interface (Figure 1) and it is available as open source code.

**Figure 1:**
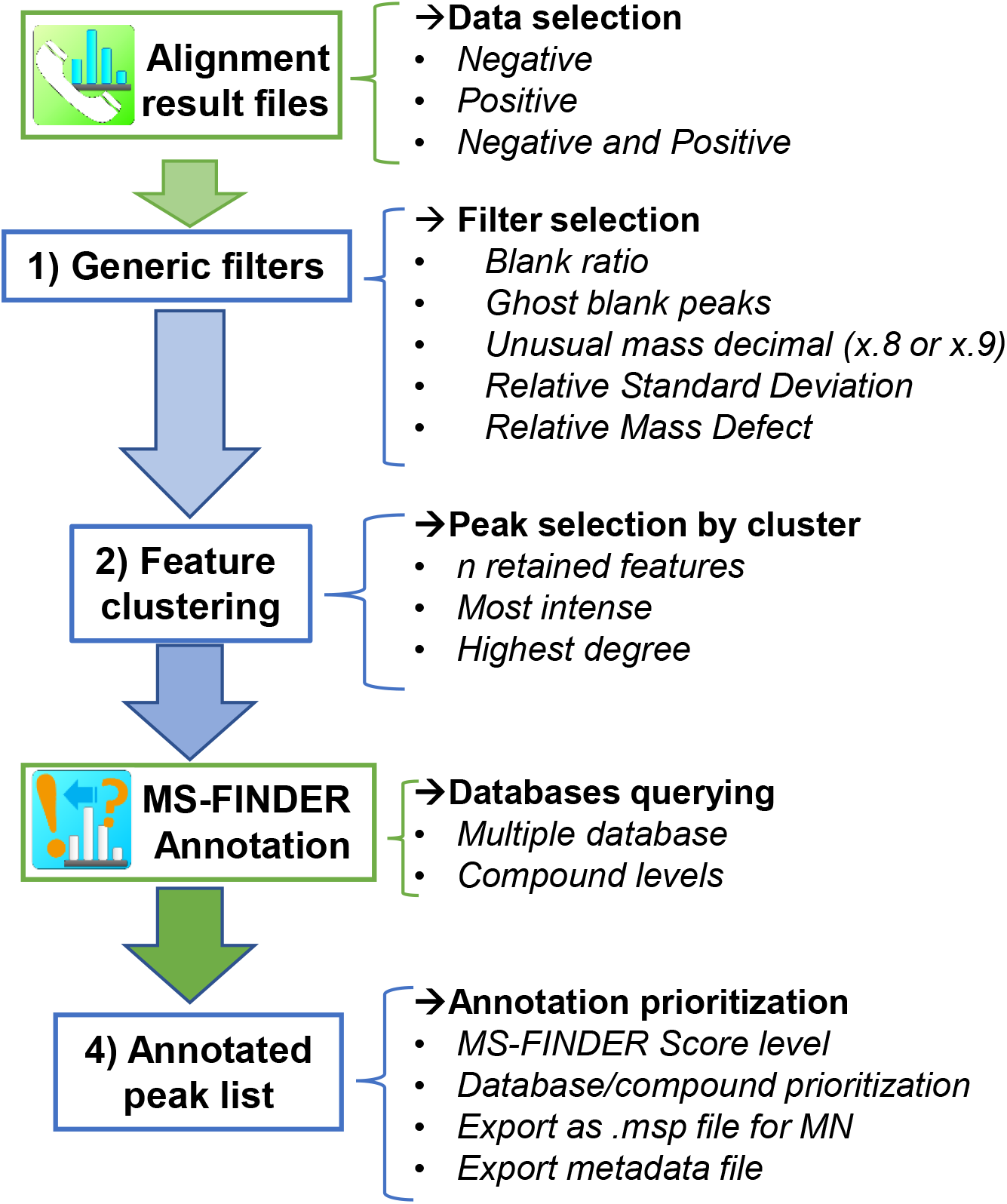
MS-CleanR workflow. Description of each step in the Shiny user interface workflow and the options available at each step. (MN, mass spectral similarity networking)

## METHODS

### Standards

Individual solutions of natural products (NPs) compounds (Metasci, Toronto, Canada) were prepared at 100 μg/mL in H_2_O or MeOH according to the supplier’s recommendations. Mixes of 10 compounds were prepared by pooling 10 μL of each individual solution to a final concentration of 10 μg/mL. We selected 51 NPs eluting from 2-18 minutes as a first test mixture to construct DB-level 1 annotation.

IROA Mass Spectrometry Library of Standards (Sigma-Aldrich, Darmstadt, Germany) in 96-well plate format (5 μg per well) were used. The contents of each well were dissolved in 50 μL of solvent (5% MeOH or MeOH/CH_3_Cl/H_2_O 1:1:0.3), as recommended by the manufacturer, to obtain a concentration of 100 μg/mL. Each plate was then sonicated for 5 minutes. Mixes of up to 12 compounds with distinct exact masses were obtained by pooling 20 μL from each well. The final concentration in each mix was 8 μg/mL. We selected 167 standards eluting from 2-18 minutes as a second test mixture.

### Plant material

*A. thaliana* (wild-type Col-O) were grown either in hydroponic culture, in plastic pots (high density), or in Jiffy® pots. For hydroponic culture, seeds were sown in 96 plates in MS liquid medium + 1% sucrose. After 11 days, the medium was replaced by MS medium. After 14 days, seedlings were collected and gently dried on absorbent paper. For culture in plastic pots, seeds were sown densely on soil in plastic pots and cultivated in a growth chamber with a cycle of 16h light-8h dark, at 22°C in the light and 20°C in the dark, and at 80% relative humidity. After 21 days, the aerial parts of the plants were collected. For culture in Jiffy® pots, three seeds were sown per pot and cultivated in a growth chamber, as for the plants in plastic pots. After 32 days, rosette leaves were harvested. For each growing condition 200 mg of plant material per sample were collected, placed in a FastPrep tube (MP Biomedicals Lysing Matrix D, Illkirch, France) and frozen in liquid nitrogen. For extraction, each sample was ground with a Mixer Mill MM 400 grinder (Retsh, Eragny sur Oise, France) by applying two cycles of 30 seconds at 30 m/sec. Biphasic sample extraction was adapted from Salem *et al*. 2016^23^ by adding two cycles of 20 seconds at 6 m/sec. in the FastPrep-24™ benchtop homogenizer (MP Biomedicals™, Illkirch, France) in 1 mL M1 (methyl tert-butyl ether/methanol, 3:1, v:v) extraction solution. After grinding, FastPrep tube was transfered in glass tube and 5.7 mL of M1 was added with 4.3 mL of M2 (water:methanol, 3:1, v:v) and vortexed for 1 min. The phases were separated by centrifugation at 4°C and 4000 rpm for 5 minutes. The aqueous phases (400 μL) were evaporated under nitrogen and the extracts were resuspended in 750 μL MeOH:H_2_O (1:1). Samples were filtered through 0.2 μm PTFE filters (Thermo Scientific™) and transferred to vials. An extraction blank (without plant material) and a QC (Quality Control) sample (aliquot of all samples) were also prepared to validate the LC-MS profiles.

Seeds of *Medicago truncatula* strains A17, DZA45.5 and F83005.5 (called F83 hereafter) were scarified with sand paper, sterilized in 3.2% bleach for 2 min then rinsed in water four times before soaking in water for 20 min. Seeds were placed on water agar and placed at 4°C for 4 days then for one night at 25°C to germinate. Germinated seedlings were transferred onto M medium^24^ then placed in a growth chamber at 22°C and 50% humidity with cycles of 16h light-8h dark for 14 days. The roots were ground with a Mixer Mill MM 400 grinder by applying two cycles of 30 seconds at 300 Hz. One hundred milligrams of ground tissue were placed in 2 mL FastPrep tubes containing 1.4 mm ceramic spheres (Lysing Matrix D) and extracted with 1 mL of acidified aqueous solution of methanol (MeOH/H_2_O/HCOOH, 80:19:1). After two cycles of 20 seconds at 6 m/sec. in the FastPrep-24™ (MP BiomedicalsTM), the samples were centrifuged at 4°C and 10 000 rpm for 10 minutes. The supernatants were transferred into vials. An extraction blank and QC (Quality Control) were also done for extraction and analytical validation.

### UHPLC-HRMS profiling

Ultra High Performance Liquid Chromatography-High Resolution MS (UHPLC-HRMS) analyses were performed on a Q Exactive Plus quadrupole mass spectrometer, equipped with a heated electrospray probe (HESI II) coupled to an U-HPLC Ultimate 3000 RSLC system (Thermo Fisher Scientific, Hemel Hempstead, UK). Samples were separated on a Luna Omega Polar C18 column (150×2.1mm i.d., 1.6μm, Phenomenex, Sartrouville, France) equipped with a guard column. The mobile phase A (MPA) was water with 0.05% formic acid (FA) and mobile phase B (MPB) was acetonitrile with 0.05% FA. The solvent gradient was: 0 min, 100% MPA; 1 min 100% MPA; 22 min, 100% MPB; 25 min, 100% MPB, 25.5 min, 100% MPA; 28 min, 100% MPA. The flow rate was 0.3 mL/min, the column temperature was set to 40°C, the autosampler temperature was set to 10°C and injection volume fixed to 2 μL for standard mixes and plant extracts. Mass detection was performed in positive ionization (PI) and negative ionization (NI) modes at 30 000 resolving power [full width at half maximum (FWHM) at 400 m/z] for MS1 and 17 500 for MS2 with an automatic gain control (AGC) target of 1e5. Ionization spray voltages were set to 3.5 kV (for PI) and 2.5 kV (for NI) and the capillary temperature was set to 256°C for both modes. The mass scanning range was *m/z* 70-1050 Da for standards and *m/z* 100-1500 Da for plant extracts. Each full MS scan was followed by data-dependent acquisition of MS/MS data for the six most intense ions.

### Data processing

LC-MS data were first processed with MS-DIAL version 4.12. MS1 and MS2 tolerances were set to and 0.05 Da, respectively, in centroid mode for each dataset. Peaks were aligned on a quality control (QC) reference file with a RT tolerance of 0.1 min and a mass tolerance of 0.015 Da. Minimum peak height was set to 70% below the observed total ion chromatogram (TIC) baseline for a blank injection. MS-DIAL data was cleaned with MS-CleanR by selecting all filters with a minimum blank ratio set to 0.8, a maximum relative standard deviation (RSD) set to 30 and a relative mass defect (RMD) ranging from 50-3 000. The maximum mass difference for feature relationships detection was set to 0.005 Da and maximum RT difference was set to 0.025 min. The Pearson correlation links were considered only for biological datasets with correlation ≥0.8 and statistically significant α = 0.05. Two peaks were kept in each cluster: the most intense and the most connected. The kept features were annotated with MS-FINDER version 3.26. The MS1 and MS2 tolerances were set to 5 and 15 ppm, respectively. Formula finder were exclusively processed with C, H, O, N, P and S atoms. DBs based on the genus and the family of the plant species (Table S3, Table S4, Table S7, Table S8) being investigated were constituted with the dictionary of natural product (DNP-CRC press, DNP on DVD v. 28.2) and the internal generic databases used were KNApSAcK, PlantCyc, HMDB, LipidMaps, NANPDB and UNPD. Annotation prioritization was done by ranking genus DB followed by Family DB and then generic DB (internal DB from MS-FINDER).

### Statistical analysis

Statistical analyses were done by using SIMCA-P+ (version 15.0.2, Umerics, Umea, Sweden). All data were scaled by unit variance (UV) scaling before multivariate analysis. The orthogonal projection to latent structure using discriminant analysis (OPLS-DA) was used to separate data according to *A. thaliana* growing conditions. The OPLS regression model used for the *Medicago* datasets was tuned with line resistance as the Y input: the following resistance indices 0, 1 and 2 were respectively indicated for the F83, A17 and DZA45.5 strains. Coefficient scores were used to rank variables according to their class biomarker: a high coefficient indicating a strong correlation with resistance traits.

### Mass spectral similarity network

The .msp NI and metadata files generated at the end of the MS-CleanR workflow were imported into MetGem^25^ (version 1.2.2). A mass spectral similarity network was created with a cosine score cut off fixed at 0.65, a maximum of ten connections between nodes and at least four matched peaks. The resulting network was then imported into Cytoscape^26^ (version 3.7.2) to tune visualization. Nodes were thus colored according to their annotated chemical classes and their sizes were indicated relative to the OPLS coefficient score. Edge width was deepened according to their cosine value.

## RESULTS AND DISCUSSION

### MS-CleanR Workflow and Implementation

#### Step 1: generic filters

We first applied several generic filters to pre-clean the feature table of noise. Starting from the alignment result file exported from MS-DIAL, the ratio between the mean of blank samples and quality control (QC) samples (pool of all extracts) was calculated. All features exceeding the user-defined threshold for this ‘blank ratio’ were removed. The ratio was calculated by using the height of each feature because the normalized height can produce an increase in some blank signals. This filter is also available in MS-DIAL, but MS-CleanR provides additional options for filtering ion features. A second filter, called ‘ghost blank peaks’, is based on the high background ion drift we observed in blank injections and in other samples that had a significant retention time (RT) shift (Figure S1). These peaks had a low ratio of blank to sample class that excluded them from the usual blank filtering approach. A third generic filter is based on an ‘unusual mass decimal’. When singly charged ions of basic organic molecules containing carbon, hydrogen, oxygen, and nitrogen are considered, ion features with a value of more than eight at the first decimal place of *m/z* (mass to charge ratio) are generally considered to be artifacts: this filter option can be disabled when working with exceptions (e.g., phosphorylated compounds). A fourth generic filtering approach is the ‘relative standard deviation’ (RSD) among sample classes. A high RSD value highlights poor ionization repeatability. In our implementation, the RSD value was computed for each sample class and features were removed if the RSD values in all sample classes exceeded a user-defined threshold. This approach avoids incorrect feature deletions: in the case of large sample cohorts, for example, repeated QC injections usually result in large RSDs because of a high dilution effect in the samples. Finally, we introduced a fifth filter based on the ‘relative mass defect’ (RMD) calculation. The RMD is calculated in ppm as [(mass defect/measured monoisotopic mass) × 10^6^)]. It can be used to filter compound classes^23^ and it should also be useful to remove artifactual signals. Based on all compounds exported from the Dictionary of Natural Products (DNP; available on DVD v.28.2 from CRC Press), we found that 95% of natural products (NP)s had RMD values of 156.5-969.6 ppm. When this window was extended to 99% of NPs, the range of RMD values was 52.05-2902.9 ppm.

#### Step 2: Feature clustering

To improve further the filtering process, we implemented a features clustering function to be applied to those features remaining after the generic-based filtering described above. The main goal of this step is to select the features arising from a unique metabolite signal among each cluster by using the multi-level optimization of modularity algorithm^28^. Feature clustering is first based on the peak character estimation algorithm computed by MS-DIAL, which aggregates several possible relationships at the same RT range: ion correlation among samples, MS/MS fragments in higher *m/z*, possible adducts and chromatogram correlations^22^. Additionally, we also implemented an index of possible neutral loss and a calculation of dimers/heteromers to tag clustered feature relationships. Optionally, Pearson’s correlation between features located in the same RT window (typically of 0.025 minutes) can be computed, the strong correlation links being then considered during the clustering process. If the study involves the same set of samples acquired both in PI and NI mode, the MScombine^29^ tool, incorporated into MS-CleanR, can be used to detect possible links between positive and negative features appearing in the same RT window. This process corrects misidentified relationships to consider observed *m/z* differences acquired between both ion modes. The package can only treat PI or NI data independently, however. We observed that a unique metabolite signal in each cluster can be selected by: a PI/NI adduct link (e.g. [M+H]^+^/[M−H]^−^, [M+Na]^+^/[M+FA−H]^−^; the most intense peak of the cluster, and the peak with the most relationships to other features (i.e. the highest ‘degree’ of connection). Among each cluster, one to *n* features (tunable by the user) can be selected for further annotation: the most intense, the most connected or both. The other features are removed from consideration.

#### Step 3: Feature annotation

After the above filtering steps, only a portion of the original features are exported to MS-FINDER, which greatly accelerate the processing time. This software computes feature annotations by querying internal DBs or imported DBs. Several DBs can be used to annotate a single set of features by exporting the results for each DB used. Additionally, a “compound level” column can be added into external DBs to further prioritize annotation within each DB used.

#### Step 4: Annotated peak list

This final step selects for each feature the best annotation among match possibilities exported from MS-FINDER. In the case of multiple DB interrogation, the workflow allows compound annotations to be ranked based on MS-FINDER score only or by prioritizing certain DBs, depending on user choices. This latter function can greatly improve the annotation accuracy particularly when dealing with taxonomically defined extracts^30^. MS-CleanR can also prioritize compounds based on “Compound_level” column tuned by the user in external DBs used for MS-FINDER annotation. Finally, the resulting annotated peaks list can be converted into an .msp format for mass spectral similarity networking as in GNPS^31^ or MetGem^25^ (for the detailed mathematics of the workflow, see Supporting Information Text 1).

### Workflow benchmarking on pooled standards

To validate our approach, we benchmarked the MS-CleanR workflow by using a mixture of 51 NPs standards profiled in NI and PI modes with a reverse phase column and a 25 minutes gradient. The resulting data were compiled in an in-house DB comprising RT, HRMS and MS/MS fragmentation patterns (DB-level 1 annotation according to the Metabolomics Standards Initiative-MSI^32^). To test whether the workflow retained features arising from unique metabolites and removed useless signals, we compared the results obtained by using a combination of MS-DIAL and MS-FINDER and DB-level 1 annotation to those obtained by using MS-CleanR. For the latter, we created another DB of the same metabolite set encompassing accurate mass, molecular formula and SMILES strings (DB-level 2 annotation according to the MSI) to reproduce real-case annotation processing. All five generic filters were used and the two most intense and two most connected features within each cluster were exported for annotation by using the ‘formula prediction and structure elucidation by *in silico* fragmentation tool’ in MS-FINDER (Table S1).

As anticipated, we observed significant feature inflation in this mixture of 51 NP standards: 869 signals from PI and NI acquisition modes were detected (Figure 2). This approximately 95% feature inflation is consistent with a previous report of 10 000-30 000 features detected after injection of 900 unique metabolites^33^ and with a study that used isotope labeling as a feature filtering approach^11^. Blank ratio filtering deleted 50% of the features and the other generic filters described above removed 15% of the remaining ones. Feature clustering resulted in a further reduction of 18%, resulting in a total of 115 features retained. Overall, the workflow filtered out 80% of all detected signals. By using this approach, there was a remarkable improvement in the annotation rate (unique metabolites/detected features) from 5% to 45% (Figure 2). Consequently, 21 metabolites displayed an isolated *mz*-RT signal whereas the others were grouped in clusters of two to eleven features (Figure 3A). Overall, 50 metabolites were annotated, 44 of which matched perfectly with level 1 annotation DB (Table S1). The remaining ones were annotated as an isobaric/isomeric match because of prioritization of highest MS-FINDER scoring value (e.g., 4-Aminosalicylic acid and 5-Aminosalicylic acid). In the case of gramine, for example, the major pseudo-molecular ion had an *m/z* value of 130.06493 at RT 7.75 minutes (Figure 3B). By applying feature clustering, we detected an in-source fragment corresponding to the neutral loss of the dimethylamine group at *m/z* 130.0649. This feature was removed and only the signal at *m/z* 103.054 and *m/z* 175.1228 were exported for annotation. Since *m/z* 175.1228 was the most intense peak, it was retained and annotated as gramine (Δppm=0.4) with a perfect match. The peak detected at RT 11.47 minutes was grouped in a cluster of 11 features, mainly related to similar MS/MS spectra. In this case, the PI and NI clusters were merged according to their detected adduct ([M+H]^+^ and [M−H]^−^, respectively) and the feature with highest MS-Finder annotation score was retained in the final peak list and identified as neohesperidin dihydrochalcone (Δppm=0.4). Formononetin displayed complex adduct relationships in PI and NI modes and successive features with higher *mz*’s MS/MS fragment of formononetin in PI mode. The merging of PI and NI modes allowed the main feature in this complex cluster to be selected and provided a perfect match with level 1 annotation DB. The only mismatch was encountered for phloridzin due to the neutral loss of a glucose moiety in both in PI and NI modes. Only genine was detected in PI mode, resulting in selection of this signal in the final peak list.

**Figure 2.**
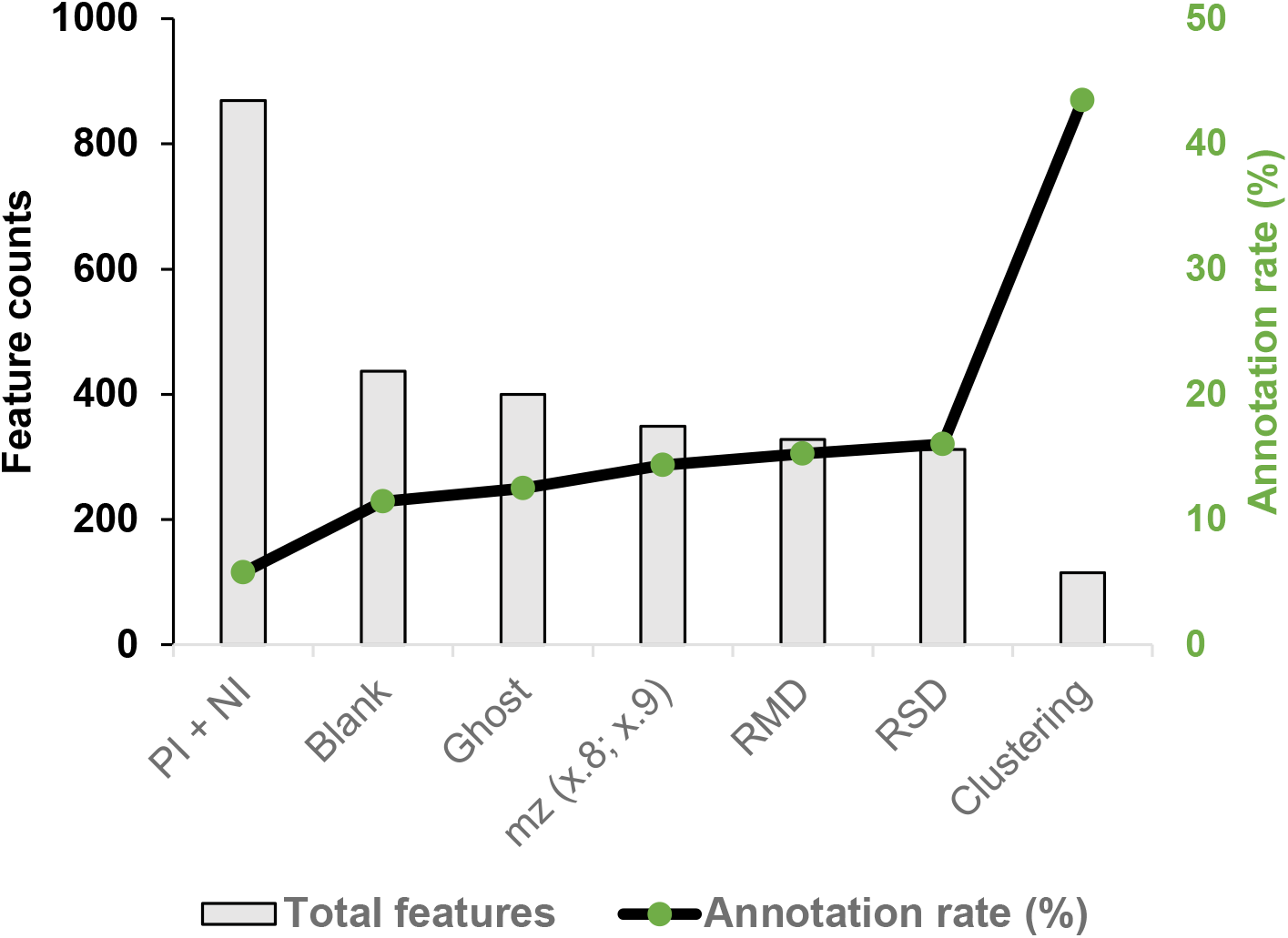
Feature filtering of the LC–MS dataset from 51 NPs standards. Generic filters and the feature clustering algorithm were applied to the initial PI + NI mode dataset. The bar plot displays feature counts after successive filters. The line plot displays annotation rate (unique metabolites/feature counts in %).

**Figure 3.**
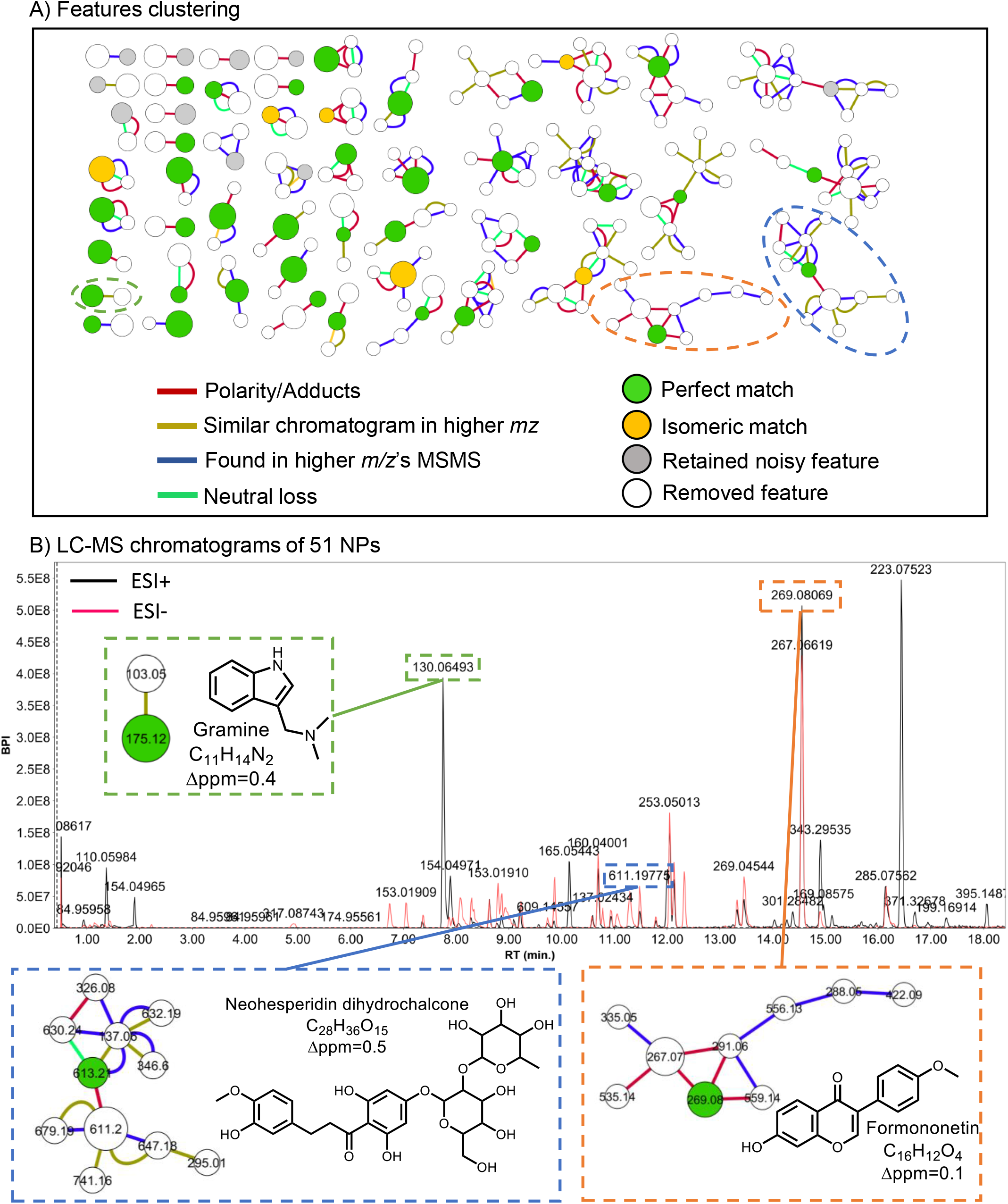
MS-CleanR feature clustering of 51 NPs. Clustering was based on the peak character estimation and multi-level optimization of modularity algorithms. A) Cluster plot of the whole dataset excluding size one clusters. B) UHPLC-HRMS base peak intensity (BPI) chromatogram of the standards mixture containing 51 NPs. Three representative compounds and their respective clusters are indicated.

To model more closely a real biological sample, we standardized our workflow by using a mixture of 167 standard compounds from the IROA Mass Spectrometry library (Table S2). As above, we found significant feature inflation: 6732 signals after concatenation of PI and NI datasets (Figure S2). Unlike the standardization with NPs, above, the generic filters removed only 15% of features. The most important improvement was obtained by feature clustering, which filtered out 90% of the detected features leaving 611 signals. Among these, 127 features were identified with a perfect match compared to Level-1 annotation DB and 21 were annotated as an isomeric match (Table S2). Twelve features were removed due to their co-elution with other compounds and four had a significant RT shift due to their poor peak shapes. The final three compounds were not annotated because of neutral loss of the same moiety in PI and NI modes, which led to their misidentification. Overall, the annotation rate with this workflow was 27% (Figure S2) and 90% of unique metabolites were retained.

### Evaluation of MS-CleanR on biological samples

To evaluate the utility of the workflow on a real dataset, we set up an experiment to compare metabolome changes in *Arabidopsis thaliana* plants due to different culture conditions and age of the plants. Three cultural conditions were assessed (low density growth in Jiffy® pots for 32 days, high density growth in plastic pots for 21 days and hydroponic culture in liquid MS medium for 14 days) and 10 biological replicates were analyzed per culture condition. At harvest time, 4 leaves (2 cotyledons and 2 leaves) were observed for hydroponic plants, the densely seeding plants showed not more than two small, but completed, developed leaves, while the jiffy growing plants harbored large and well developed rosette leaves. Extracts were made from the aerial parts of the plants grown in pots and from the roots and green tissues of plants in hydroponic culture, and the extracts were profiled by LC-MS. The datasets acquired in PI and NI modes were treated by using the MS-CleanR workflow with default parameters (see Methods). Sequential principal component analysis (PCA) was used to provide an unsupervised overview of the LC-MS fingerprints resulting from the generic filters and feature clustering (Figure 4). The PCA score plot of raw PI and NI mode data displayed 51% of total explained variance using the first two principal components. QC samples appeared in the center of the PCA score plot, demonstrating the reproducibility of the LC-MS analysis. As expected, the youngest plants growing hydroponically were completely separate on the first principal component (PC1) axis from the older plants growing in pots. The plants growing in Jiffy pots and plastic pots could not be distinguished in the raw dataset. After the generic filter step, the data from these latter two conditions formed more distinct clusters, the total explained variance was slightly improved at 58% and the number of features decreased by 35% (Figure 4). After the feature clustering step, the number of features was reduced by 80%. All datasets were annotated with in-lab DB (level 1) and with MS-FINDER (level 2) by reference to external DBs of *Arabidopsis* (Table S3) and Brassicaceae compounds (Table S4) and an internal MS-FINDER plant-related DB (comprising PlantCyc, KNApSAcK, HMDB, LIPID MAPS and UNPD). In the raw PI and NI dataset exported from MS-DIAL (1163 features), 42% of all features were annotated, 26% of them appeared in the *Arabidopsis* DB, 2% in the Brassicaceae DB,7% in the internal MS-FINDER DBs and 6% with in-lab DB (Figure 4); 58% of all features were unidentified. The generic filters removed 15% of all features and increased the annotation rate to 59%. Feature clustering drastically reduced the number of features (254 *m/z* × RT pairs) and increased the annotation rate to 73%. Using annotation DB prioritization, 53% of retained features were annotated in *Arabidopsis* genus and 13% at level 1 with in-lab DB, only 27% remained unidentified. Orthogonal projections to latent structures discriminant analysis (OPLS-DA) of the most highly ranked features identified three amino acids (oxoproline, citrulline and glutamine) that discriminate between growth in pots and hydroponic growth (Table S5). This may be related to differences in nitrogen availability in the hydroponics medium and in potting soil.

**Figure 4.**
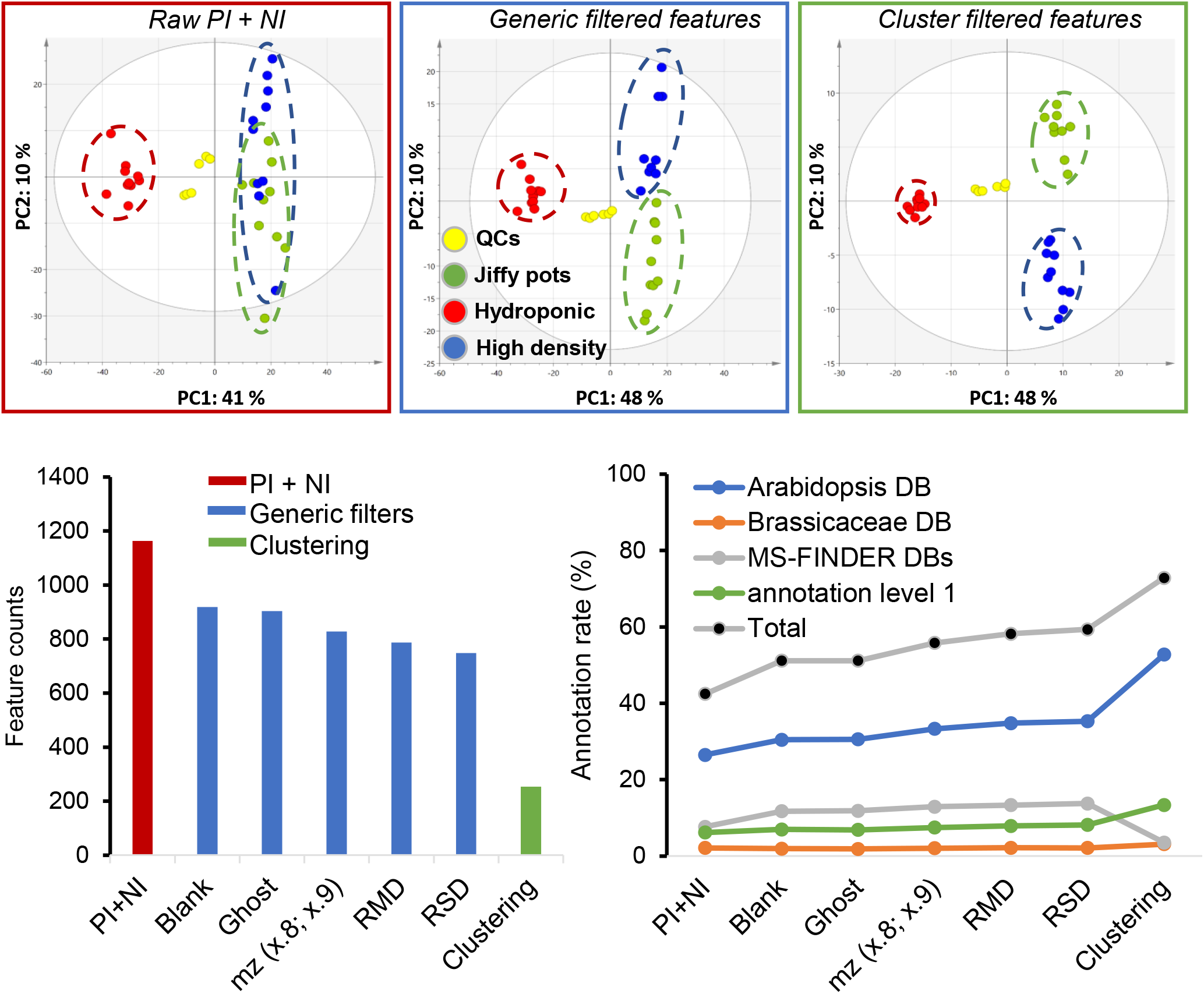
LC–MS dataset processing of the metabolomes of *A. thaliana* plants growing in different conditions. Top: Sequential PCA score plots of raw PI and NI mode data and the data after applying generic filters and feature clustering. Dotted circles indicate biological sample type distribution (yellow, QC injections; green, plants growing in Jiffy pots at low density; blue, plants growing in plastic pots at high density; red plants in hydroponic culture). Bottom: The bar plot shows the feature counts after successive filtering steps. The line plot displays the annotation rate (unique metabolites/feature counts expressed as %) after successive filtering steps using annotation DBs prioritization.

### Metabolic profiling with MS-CleanR

Untargeted metabolomic profiling has emerged as a method of choice to identify metabolic markers associated with beneficial traits in plants, such as resistance to biotic stresses. In this context, the MS-CleanR workflow could greatly improve the results of untargeted metabolomics. To illustrate this point, we used as a model the legume *Medicago truncatula* and the pathogenic oomycete *Aphanomyces euteiches*, a major pathogen of several legume species^34^. Genome-wide association studies of 179 lines of *M. truncatula* have identified major loci involved in the resistance of the plant to *A. euteiches*. Moreover, genes encoding enzymes involved in the synthesis of antimicrobial metabolites are expressed in uninfected plants^35^. This suggests that antimicrobial metabolites in uninfected plants may be useful biomarkers with which to select legumes lines resistant to *A. euteiches*. To identify these metabolites, we applied the MS-CleanR workflow to analyze the metabolomes of roots from three different strains of *M. truncatula* that have different levels of resistance to *A. euteiches* infection: strain DZA45.5 has the highest level of resistance, A17 an intermediate level, and F83 is the most susceptible^36^. These three strains were analyzed by LC-MS in NI mode and potential biomarkers were highlighted by multivariate data analysis (Table S6). The metabolites that were differentially produced in the two most resistant strains (A17 and DZA45.5) when compared to the more sensitive one (F83) were identified by OPLS regression.

After application of the MS-CleanR workflow, the PCA score plot showed a net clustering of the samples from each strain of *M. truncatula*. QC samples were centered on the PCA plot demonstrating very good reproducibility (Figure 5). When annotated by reference to DBs from *Medicago* or the legume family Fabaceae, 60% of the dataset was annotated (Figure 5) and an additional 9% with MS-FINDER DBs. A molecular spectral similarity network was built to highlight common chemical class related to resistance traits (Figure 6). Among all annotated features, flavonoids and terpene glycosides compounds were prevalent. This latter class encompass mostly triterpene sapogenins which appeared to be highly correlated to the resistance traits according to the OPLS regression model. In particular, the four top ranked compounds belonged to two clusters related to sapogenins and one to flavonoids. Our untargeted approach revealed the presence of Apigenin-7-O-glucuronopyranoside (best MS-FINDER score among several possible match in flavonoid class) only in the resistant DZ45.5 strain. This result corroborated a previous study by our group which demonstrated the implication of flavonoid pathway in resistance^35^. However, other detected flavonoids were not correlated to the resistance contrary to sapogenins class. Among the 151 terpene glycosides annotated in this study, 36 were also identified by a large-scale sapogenin profiling study in various ecotypes of *M. truncaltula*^37^ (Table S6). Interestingly, the three-top ranked sapogenins by OPLS model (Azukisaponin III, Arjunolic acid 3-glucoside and Soyasaponin I) displayed an isobaric match with tow hederagenin glycoside and a bayogenin derivatives respectively annotated by Sumner and colleagues. These sapogenins accumulates preferentially in roots than in leaves. These organs, however, have distinct profiles of specific saponins, which may be explained by the adaptation of each ecotype to its biotic environment. A previous study, for example, showed that saponins derived from hederagenin glycoside in *M. truncatula* have antifungal activity^38^. Our study confirmed a higher level of these compounds in the strains resistant to *A. euteiches* (DZA45.5 and A17) than it is in the sensitive strain F83. Although the relevance of saponins to resistance of *M. truncatula* to *A. euteiches* remains to be confirmed, these findings demonstrate the potential value of applying metabolomics tools to identify biomarkers of plant resistance.

**Figure 5.**
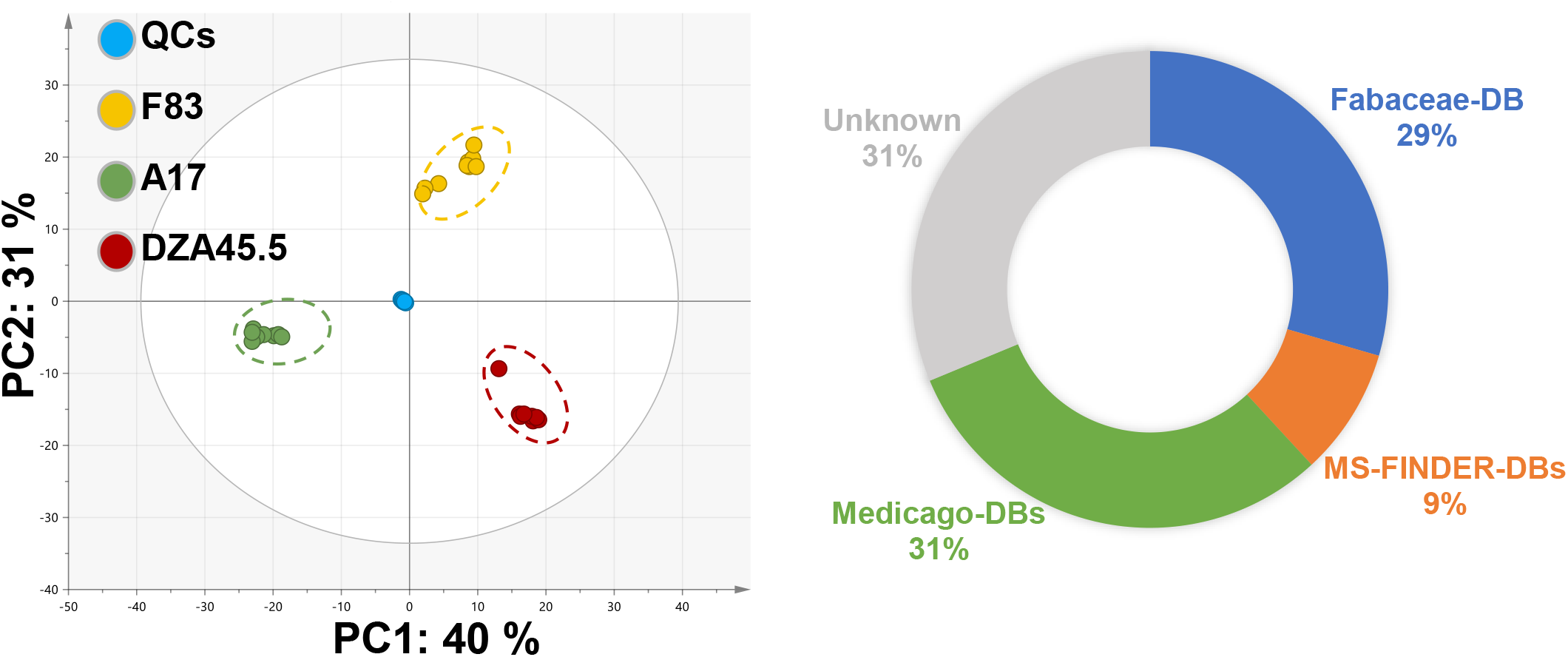
LC–MS NI dataset processing of the metabolome of roots from three strains of *M. truncatula.* Left: PCA score plot after applying the MS-CleanR workflow. Dotted circles enclose samples from each plant strain. Right: Circular plot of the proportions of features annotated with reference to the indicated databases (DB).

**Figure 6.**
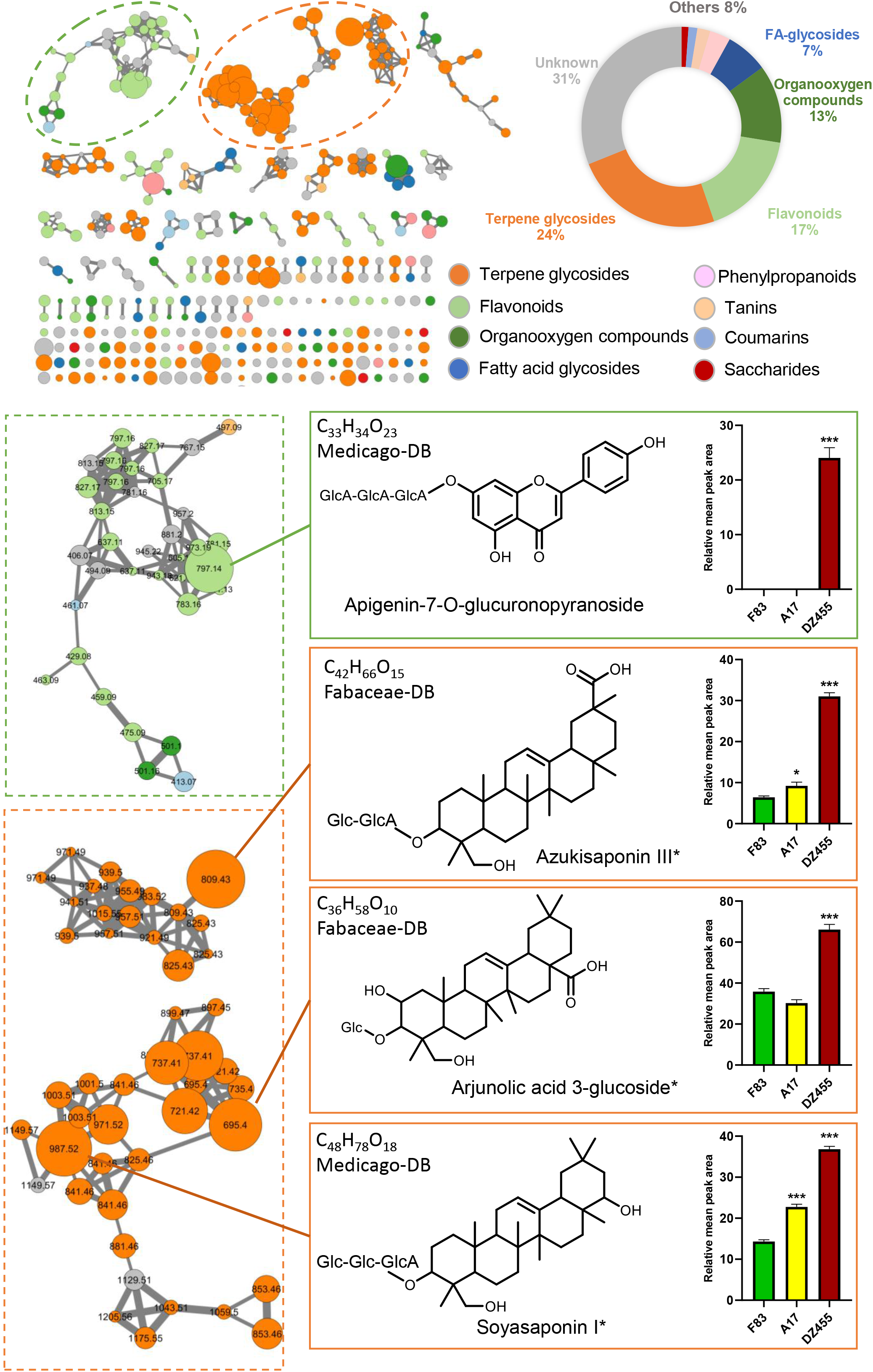
Mass spectral similarity network of *M. truncatula* NI dataset (cosine ≥ 0.8). Nodes are colored according to their chemical classes and sized relative to their OPLS regression coefficient score (See text for details). Edge width is proportional to cosine value. Pie chart display annotated chemical class ratio in LC-MS NI dataset (Others include coumarins derivatives, tanins and saccharides chemical classes). Bar plots display normalized mean peak areas for the four most highly ranked structures by OPLS-regression modeling (Table S6). One-way ANOVA and Dunnett’s post-hoc test (p≤0.05) were used to assess differences between the sensitive (F83) and resistant (A17 and DZA45.5) *M. truncatula* strains (p≤0.05: *; p≤0.01: **; p≤0.001: ***). Compound names with asterisk indicate an isobaric annotation match with ref 37. (Glc: Glucoside, GlcA: Glucuronopyranoside)

## CONCLUSIONS

The main goal of LC-MS-based untargeted metabolomics is to convert chromatographic profiles of complex biological extracts into a comprehensive metabolite list. Professor Ian Wilson summarized the challenge thus: “LC-MS includes everything, which means you see everything. Thus, the challenge is to take oceans of data, and make rivers of information, and finally puddles of knowledge.” (NIH Metabolomics symposium, August 2013). We demonstrate here that feature degeneracy - the ocean of data - has a great impact on the final annotated peak list information, thus impacting the biological knowledge mined from untargeted metabolomic studies. We estimate, based on analysis of standard mixtures, that feature inflation is close to 95%, in agreement with other studies^33,11^. Our package MS-CleanR, with its a point-and-click software on a Shiny interface, is a new component in the suite of tools comprising the GUI software MS-DIAL and the annotation capabilities of MS-FINDER which altogether provide a comprehensive workflow, from raw data to final annotated peaklist. MS-CleanR can reduce the number of features by 80-90% and keep most unique metabolite signals without compromising the final data structure. The opportunity to rank the annotation results with reference to in-house databases narrows down the final identification possibilities. Additionally, the package is able to combine both PI and NI mode (*A. thaliana* experiment) or to treat only one mode (*M. truncatula* study) depending of the study objectives. We demonstrate the utility of this workflow by analyzing secondary metabolites levels in three *M. truncatula* strains with different susceptibilities to a pathogenic oomycete. We could annotate 70% of the dataset with 60% at the genus or family level using DBs prioritization. The resulting mass spectral similarity network further supports annotation results since most clusters gathered the same metabolite chemical class. Still, our approach was unable to keep only unique metabolite features regarding the annotation rate comprising between 24 and 45% for standard mixtures. A limitation of our filtering process is its dependence to chromatographic resolution, which can seriously impair the final results by clustering several unique metabolites together. In the present study, we chose a twenty minutes gradient, like those generally applied in most untargeted metabolomics studies. Extending the elution time might improve the chromatographic resolution but is difficult to apply in day-to-day work, especially for high-throughput experiments. These challenges will be addressed in future developments of MS-CleanR.

## ASSOCIATED CONTENT

The Supporting Information is available free of charge on…

- figure S1. Alignment spot screenshot in ESI PI and NI ionization modes showing repeated blank pseudomolecular ions detected massively in QCs samples with a retention time shift. (PDF)
- Figure S2. Features filtering of LC-MS dataset from 167 IROA-MS standards library according to generic filters and clustering algorithm. Barplot display feature counts after successive filters. Line plot display annotation rate (unique metabolites/feature counts in %). (PDF)
- Sup Table S1: Excel table with cluster annotation, result summary and database for level 2 annotation imported in MS-FINDER for the 51 NPs dataset (XLSX)
- Sup Table S2: Excel table with cluster annotation, result summary and database for level 2 annotation imported in MS-FINDER for the 167 IROA MS standard library dataset (XLSX)
- Sup Table S3: Arabidopsis DB used as input for level 2 annotation in MS-FINDER (TXT)
- Sup Table S4: Brassicaceae DB used as input for level 2 annotation in MS-FINDER (TXT)
- Sup Table S5: Arabidopsis dataset treated by MS-CleanR and OPLS-DA top ranked features (XLSX)
- Sup Table S6: Medicago dataset treated by MS-CleanR and OPLS regression coefficient for feature ranking (XLSX)
- Sup Table S7: Medicago DB used as input for level 2 annotation in MS-FINDER (TXT)
- Sup Table S8: Fabaceae DB used as input for level 2 annotation in MS-FINDER (TXT)
- Raw data from *Arabidopsis* and *Medicago* LC-MS profiling available on Zenodo using the DOI: 10.5281/zenodo.3744480

## ACKNOWLEDGMENTS

We thank Dr. Stephane Bertani for providing us standards from the IROA-MS Library. Financial support from The French national infrastructure for metabolomics and fluxomics, MetaboHUB-ANR-11-INBS-0010 and PSPC Solstice project supported by Bpifrance (SOLutionS for Integrates Treatments under Environmental Management). We thank E Amblard, N. Jariais and C. Jacquet for *M. truncatula* cultures and A. Haouy for *A. thalina* cultures and sample preparations. We also acknowledge Carol Featherstone of Plume Scientific Communication Services for professional scientific editing.

