## Supplementary material for "MS-CleanR: A feature-filtering approach to improve annotation rate in untargeted LC-MS based metabolomics": Figure S1

| | N° | RT (min.) | *mz* | Adducts | Fill % | S/N | ANOVA | FC | Blank | QCs |
|---|---|---|---|---|---|---|---|---|---|---|
| ESI- | 61 | 15.87 | 293.1762 | [M-H]- | 0.50 | 1783.1 | -1.00E00 | -1.00 | | |
| | 62 | 16.88 | 293.1763 | [M-H]- | 0.50 | 624.0 | -1.00E00 | -1.00 | | |
| | 63 | 22.85 | 293.1796 | [M-H]- | 0.50 | 3852.7 | -1.00E00 | -1.00 | | |
| | 64 | 19.78 | 297.1532 | [M-H]- | 1.00 | 421.7 | -1.00E00 | -1.00 | | |
| | 65 | 19.98 | 297.1532 | [M-H]- | 1.00 | 593.0 | -1.00E00 | -1.00 | | |
| ESI+ | 1 | 1.18 | 74.0600 | [M+H]+ | 0.50 | 344.9 | -1.00E00 | -1.00 | | |
| | 2 | 0.59 | 74.0600 | [M+H]+ | 0.50 | 58.9 | -1.00E00 | -1.00 | | |
| | 3 | 3.53 | 74.0601 | [M+H]+ | 0.50 | 47.3 | -1.00E00 | -1.00 | | |

**figure S1.** Alignment spot screenshot in ESI PI and NI ionization modes showing repeated blank pseudomolecular ions detected massively in QCs samples with a retention time shift.
