## Supplementary material for "MS-CleanR: A feature-filtering approach to improve annotation rate in untargeted LC-MS based metabolomics": Figure S2

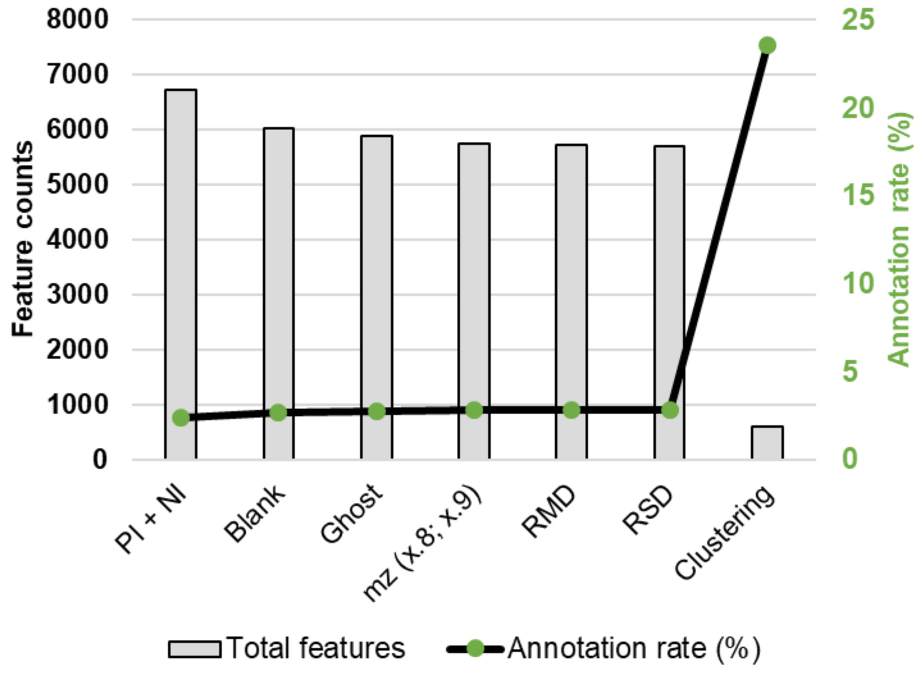


**Figure S2. Features filtering** of LC-MS dataset from 167 IROA-MS library standards according to generic filters and clustering algorithm. Barplot display feature counts after successive filters. Line plot display annotation rate (unique metabolites/feature counts in %).
