## Supporting Information Text 1 for "MS-CleanR: A feature-filtering approach to improve annotation rate in untargeted LC-MS based metabolomics"

Supplementary text from "MS-CleanR: A feature-filtering approach to improve annotation rate in untargeted LC-MS based metabolomics"

From a technical standpoint, the MS-CleanR package is organized around 4 main functions described below.

### Clean_msdial_data

This function transforms positive and/or negative MSDIAL files into an aggregated CSV file containing peaks information without noisy peaks.

First the peaks identified by MSDIAL are filtered according to various criteria:

- Filtering of blank peaks: deletion of peaks for which $\frac{avg_{Blk}}{avg_S} \geq t_{Blk}$, with $t_{Blk}$the noise threshold provided by the user, $avg_{Blk}$the average intensity of blank samples, and $avg_S$the average intensity of QC samples, or standard samples if no QC are provided, or all non blank samples if QC nor standard samples are available.
- Filtering of phantom blank peaks : Detection of high background ion drift detected in blank injections and appearing in other samples with a significative retention time (RT) shift. Based on *m/z* detected in blank samples with three digits and removed from whole feature list without taking into account retention time of blank features.
- Filtering of incorrect masses: deletion of peaks having a mass whose 1st decimal is an 8 or a 9.
- Filtering based on RSD: deletion of peaks having the RSD of all its classes greater than a given threshold $t_{RSD}$, the RSD of the samples class $X$being defined as $RSD_X = \frac{sd_X \times 100}{avg_X}$, with $sd_X$the standard deviation of class $X$' samples intensities and $avg_X$the corresponding average intensity.
- Filtering based on RMD: deletion of peaks with $RMD < t_{RMD}^{min}$ou $RMD > t_{RMD}^{max}$, with $t_{RMD}^{min}$and $t_{RMD}^{max}$thresholds provided by the user and $RMD$the relative mass defect of the peak.

The user can choose none, all, or any combinations of filters based on her needs.

The next step is to regroup peaks by clusters using the multi-level optimization of modularity algorithm [1] implemented in igraph [2]: each MS peak is considered as a node in the graph and the links taken into account are MSDIAL possible adducts links and, optionally if the user wishes so, the Pearson correlation computed on samples intensities between each pair of peaks in a given retention time window $t_{RT}$provided by the user.

Once the clusters are computed, we use MSCombine [3] to detect adducts, neutral losses and isomers in each cluster, but also between positive and negative clusters who have peaks in a $t_{RT}$retention time window.

At this point, a single MS peak can be identified as being part of several adducts relations. In order to select the best solution for all our peaks, the links are iteratively filtered:

- First a graph is created containing all possible adduct links between peaks, each peak being a node.

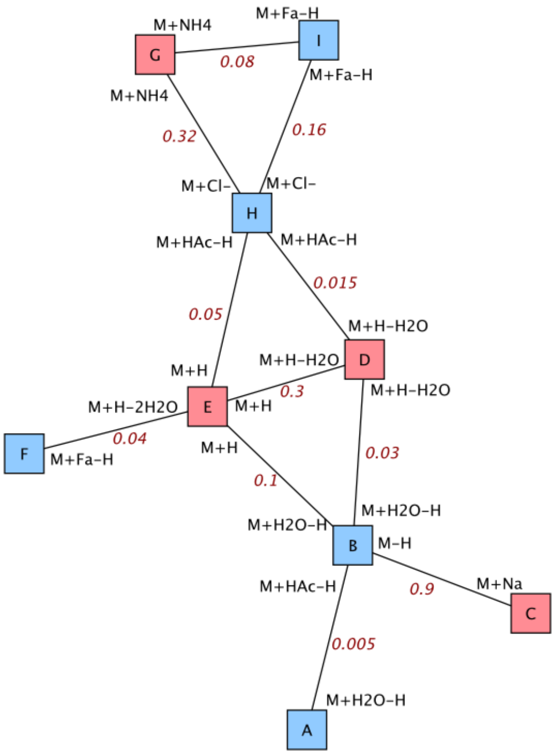


- Each edge in the graph has 3 attributes:
  - The adduct label associated with the first node (for example, [M+H]+)
  - The adduct label associated with the second node (for example, [M+Cl-]-)
  - A weight, which is the product of the probabilities of appearance for the 1st adduct and the 2nd adduct, computed on the adduct links detected by MSDIAL (if we consider that [M+H]+ has a probability of appearance of 1 and [M+Cl-]- of 0.8, the link joining 2 nodes having these labels has a strength of 0.8).

| **Adduct selection example : adducts probability of appearance** | | | |
|---|---|---|---|
| **Positive mode** | | **Negative mode** | |
| **Adduct** | **Proba** | **Adduct** | **Proba** |
| M-H | 1.00 | M+H | 1.00 |
| M+Cl- | 0.80 | M+NH4 | 0.40 |
| M+Fa-H | 0.20 | M+H-H2O | 0.30 |
| M+H2O-H | 0.10 | M+Na+ | 0.90 |
| M+HAc-H | 0.05 | M+H-2H20 | 0.20 |

- Then, while we have contradictory links in our adducts graph, we iterate on the edges:

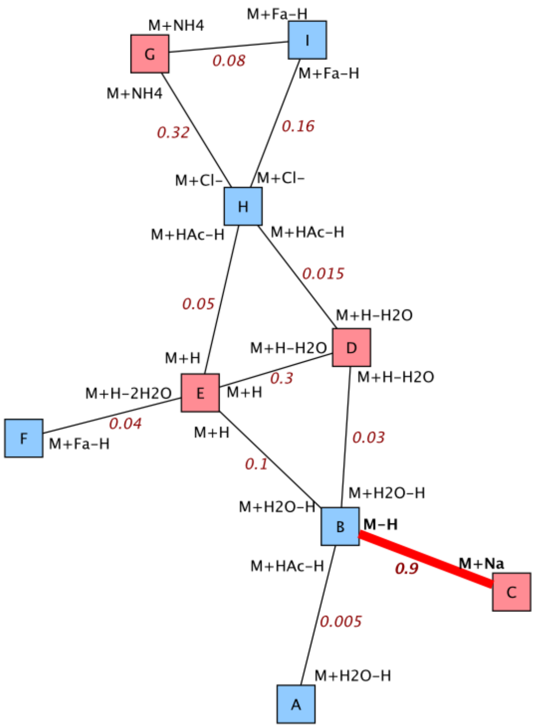


- Among the edges not yet selected, we select the edge having the highest strength $e$.

- We delete all edges contradictory to the information carried by $e$.

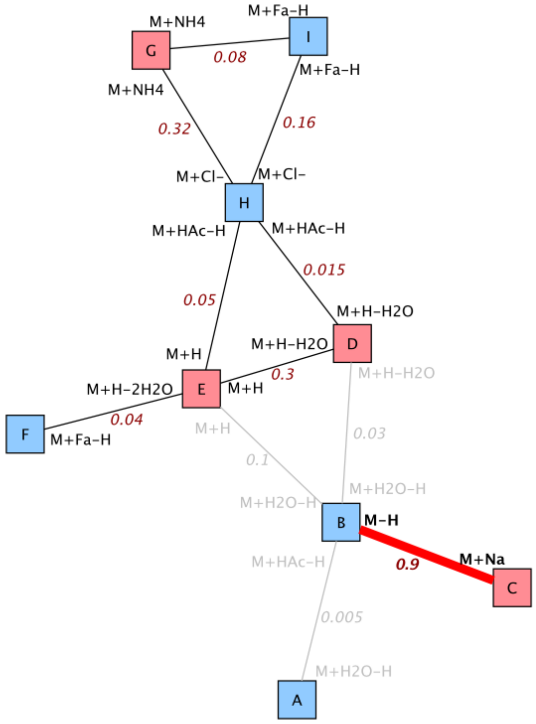


- We continue until all edges have been selected once or deleted.

We are left with a final graph free of contradictions and indicating for some of the peaks in our list the final adduct to assign them.

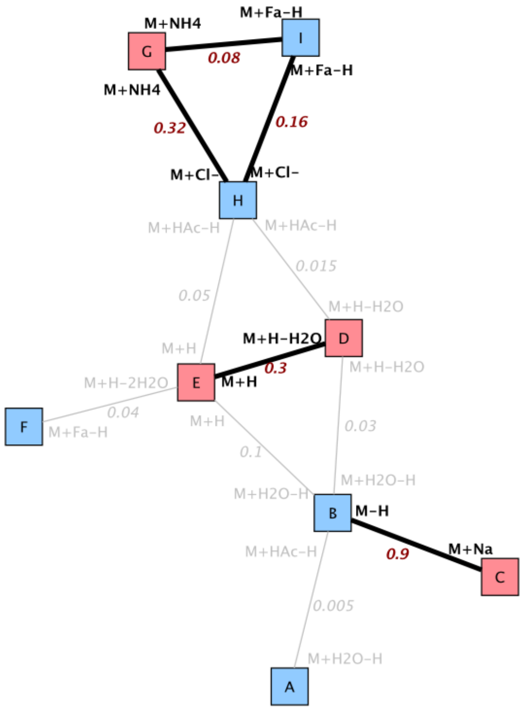

MS peaks identified as neutral losses or heteromers are deleted.
Finally we use this cleaned list of links to compute the final clusters for the MS peaks. Each cluster can contain positive peaks, negative peaks, or both.

### Keep_top_peaks

This function selects, for each cluster, a number $n$ of MS peaks from the cleaned list of peaks produced by the clean_msdial_data function. The user can choose to keep the $n$ peaks having the most links with other peaks, having the highest average intensity, or both.
It can also copy the .mat files corresponding to these peaks in a new folder for an easier import into MSFINDER.

### Launch_msfinder_annotation

This function matches the MS peaks resulting from the selection made in keep_top_peaks with possible annotations provided by MSFINDER. The user can provide a list of biosource levels $L_{bio}$ (e.g. "genus", "family", etc) and a list of compound levels $L_{cmpd}$ (e.g. "direct", "tanimoto extension", etc).

For each cluster, the function looks for the best possible annotation, it being the peak having the highest MSFINDER score in the most important possible compound level and most important possible biosource level. Compound levels are considered prior to biosource levels, and their relative importance is provided by the user: the earlier a level is in the levels lists the user gives to the function, the more important it is. For biosource levels, the last evaluated level is always the MSFINDER research in generic databases (called "generic").
For example, with $L_{bio} = (genus, family)$ and $L_{cmpd} = (direct, tanimoto\ extension)$, for each cluster the annotations will be searched within:

- First the compounds found in the genus ("direct" and "genus"),
- Then the compounds found in the family ("direct" and "family"),
- The compounds close to the genus compounds ("tanimoto extension" and "genus"),
- The compounds close to the family compounds ("tanimoto extension" and "family"),
- And finally the compounds found in generic databases ("generic").

As soon as an annotation is found for a cluster, the algorithm moves on to the next cluster. When a cluster contains 2 peaks linked by an [M+H]+/[M-H]- relation, these peaks' annotation possibilities take precedence over all others since this link is considered a strong hint to find the cluster's actual MS peak.

The final annotation result is a list of $c$ MS peaks, $c$ being the number of clusters computed in clean_msdial_data. Each cluster can have a "Simple ID" if an annotation is found, a "Double ID" if an annotation is found both is positive and negative modes, or an "Unknown Compound". For clusters without annotation, only the peak having the highest mass is kept in the final file.

Finally, for an easier reading of the final annotated peaks list, the MSFINDER annotation scores can be multiplied by personalized coefficients provided by the user for each compound and biosource level, and samples intensities are normalized.

### Convert_csv_to_msp

This function converts the annotated peaks list produced by launch_msfinder_annotation to 2 MSP files, one for positive peaks and one for negative peaks. The user can choose to export all peaks or only those having a score greater or equal to a threshold $t_{MSP}$ of her choice. It must be noted that $t_{MSP} = 0$ will not export all peaks but only those having an annotation.

### Références


1. Vincent D. Blondel, Jean-Loup Guillaume, Renaud Lambiotte, Etienne Lefebvre: Fast unfolding of communities in large networks. J. Stat. Mech. (2008) P10008
2. https://igraph.org/r/
3. https://CRAN.R-project.org/package=MScombine
